# Identification of a miRNA signature for schizophrenia in plasma-derived extracellular vesicles

**DOI:** 10.64898/2026.03.06.710021

**Authors:** Mary E W Collier, Joshua Chiappelli, Hollie Marshall, Nicolas Sylvius, Natalie Allcock, Josh Whittingham, Peter Kochunov, Robert Schwarcz, Elliot L Hong, Flaviano Giorgini

## Abstract

Background and Hypothesis: Extracellular vesicles (EVs) are phospholipid bilayer vesicles released from cells containing proteins, lipids and nucleic acids derived from the parent cell. Alterations in miRNA expression within blood-derived EVs have been proposed as potential biomarkers of disease. Specifically, identification of differentially expressed miRNAs in patients with schizophrenia (SZ) compared to healthy individuals could be used as a “miRNA signature” to aid in diagnosis and treatment. We therefore aimed to identify differentially expressed miRNAs in plasma-derived EVs between people living with SZ and healthy controls and to correlate miRNA levels with SZ-relevant clinical measures.

Study Design: Plasma-derived EVs were isolated from a cohort of 33 individuals with SZ and 34 controls. Expression of 84 miRNAs was examined using a RT-qPCR panel.

Study Results: Three miRNAs (hsa-miR-30e-5p, hsa-miR-103a-3p, hsa-miR-200b-3p) were differentially expressed between controls and patients. Gene ontology analysis of putative target genes shared between these miRNAs revealed enrichment of biological process terms related to neurogenesis. Analysis of miRNA expression compared to clinical measures showed that hsa-miR-103a-3p expression was associated with working memory and negatively correlated with white matter integrity in the combined patient-control group.

Conclusions: We have identified a miRNA signature for SZ in plasma-derived EVs and shown for the first time that hsa-miR-30e-5p expression is significantly increased in plasma-derived EVs in SZ. The genetic links between differentially expressed miRNAs and neurogenesis, along with the correlations of hsa-miR-103a-3p with working memory and white matter integrity may underlie the functional importance of altered expression of the identified miRNAs in SZ.

## Introduction

Schizophrenia (SZ) is a severe mental health condition that effects approximately 0.45 % of the adult population world-wide^1^. The main symptoms include positive symptoms such as hallucinations and delusions, negative symptoms including social withdrawal and lack of emotional expression, and cognitive impairments such as problems with memory, attention and learning. Diagnosis is limited by a lack of objective medical tests for this mental health disorder and is complicated by the overlap of symptoms with other psychiatric conditions such as bipolar disorder and major depressive disorder (MDD). Therefore, there has been a drive to explore potential biomarkers that can be used to aid in the diagnosis and stratification of patients for appropriate therapy.

Extracellular vesicles (EVs) are phospholipid bilayer vesicles that are released from all cell types, and which carry cargo derived from the parent cell such as proteins, lipids, and nucleic acids. Notably, the content of EVs can vary depending on the stimulus that induces EV release and can reflect the physiological state of the cell from which they were released^2,3^. In particular, small RNAs such as microRNAs (miRNAs) can be selectively packaged into EVs by RNA-binding proteins^4,5^. An important feature of EVs is that they protect miRNAs from degradation in the peripheral circulation. In addition, EVs are easy to isolate from blood using non-invasive techniques, making them ideal potential biomarkers. Differences in EV miRNA expression levels between healthy individuals and patients with schizophrenia therefore have the potential to be used as biomarkers to inform diagnosis, prognosis and suitable therapies^6,7,8^.

Many miRNAs are highly expressed in the brain and have important functions in the development and function of the central nervous system (CNS)^9,10^. Previous studies have shown that miRNA expression is altered in the prefrontal cortex (PFC) of patients with SZ compared to healthy controls^11,12,13,14,15^. Differences in miRNA expression between SZ patients and controls have also been detected in peripheral whole blood^16,17,18^, circulating peripheral blood mononuclear cells (PBMCs)^6,19,20,21^, and blood-derived EVs^7,8,22,23^. EVs can cross the blood brain barrier both from the periphery into the CNS and *vice versa*^24^, suggesting that EVs released from cells of the CNS such as neurons, microglia and astrocytes can enter the peripheral circulation. In addition, EVs are released from peripheral blood cells which have been shown to have altered miRNA expression in SZ^19,20,21^. A study in primates also showed that miRNA expression levels detected in the peripheral circulation correlated with miRNA expression profiles in the brain^25^, suggesting that peripheral miRNA expression may be used as an indicator of changes in the CNS during disease. The miRNA content of blood-derived EVs may therefore reflect differences in both the CNS and periphery of patients with schizophrenia. Furthermore, alterations in several miRNAs may represent a “miRNA signature” of SZ which could be used as a biomarker to aid in diagnosis and treatment strategies^6^.

A limited number of studies have examined the expression of a range of miRNAs in EVs derived from the blood of patients with SZ^7,8,22,23^. The aim of this study was to compare the miRNA content of plasma-derived EVs from patients with SZ to healthy controls using an RT-qPCR panel of 84 miRNAs typically found in plasma/serum and to identify changes in miRNA expression in schizophrenia. Expression of the three miRNAs significantly differentially expressed between controls and patients was also correlated with clinical measures relevant to SZ such as cognition and white matter integrity to examine whether miRNA expression was associated with cognitive deficits in patients.

## Materials and Methods

### Participants and clinical assessments

Participants were patients diagnosed with schizophrenia spectrum disorder (SSD) including schizophrenia, schizoaffective disorder, or schizophreniform disorder, or controls with no active psychiatric conditions. Patients were recruited from outpatient clinics and were clinically stable at time of participation. Controls were recruited from the general population through advertisements. All participants provided written informed consent. This study was approved by the Institutional Review Board of the University of Maryland Baltimore. Clinical assessments included the 20 item Brief Psychiatric Rating Scale (BPRS) to assess overall psychiatric symptoms, and the Brief Negative Symptom Scale (BNSS)^26^ to assess negative symptoms. Participants were also assessed for two cognitive measures; processing speed using the Digit Symbol Coding task of the WAIS-3^27^ in which participants use a provided key to draw symbols corresponding to numbers as fast as they can, and working memory using the Digit Sequencing task from the Brief Assessment of Cognition in Schizophrenia^28^ in which participants are asked to repeat series of digits. These were selected since deficits in processing speed and working memory are among the most robust cognitive impairments in schizophrenia^29^. At the time of study, 21 patients were taking a second generation antipsychotic, 2 were taking a first generation antipsychotic, 2 were taking both a first generation and second generation antipsychotic, and 8 were not taking an antipsychotic medication.

### Isolation of EVs from plasma

Participants were not fasting at the time of blood draw. Whole blood was collected in EDTA-containing tubes (Vacutainer), which were immediately centrifuged at 1000 *g* for 10 min.

Plasma was then aliquoted into separate tubes and stored at -80°C until assay. 71 plasma samples were processed for EV isolation in this study. Plasma samples (1 ml) were defrosted at room temperature and centrifuged at 1500 *g* for 10 min followed by 10,000 *g* for 10 min at room temperature. Exo-spin mini size exclusion chromatography columns (Cell Guidance Systems Ltd, Cambridge, UK) were equilibrated to room temperature and washed twice with 250 µl of 0.22 µm-filtered phosphate buffered saline (PBS), pH 7.4. Plasma (100 µl) was added to the top of each column and allowed to enter the column. PBS (180 µl) was then added to the top of the column resulting in the elution of EVs into 1.5 ml low protein-binding tubes. The Exo-spin columns were then washed four times with 200 µl of PBS and the process repeated until 0.5 ml of plasma had been processed. The EV fractions for each sample were pooled together and concentrated using Amicon Ultra 10 kDa molecular weight cut-off centrifugal filters (Sigma, Gillingham, UK) by centrifugation at 3000 *g* for 45 min at 4°C. EVs were recovered by inverting the centrifugal filters and centrifuging at 1000 *g* for 2 min.

### Nanoparticle Tracking Analysis

The size distributions and concentrations of plasma-derived EVs were examined by Nanoparticle Tracking Analysis (NTA) using a Nanosight LM10 instrument (Malvern Panalytical Ltd, Malvern, UK). EVs were diluted 1 in 20 in 0.1 µm-filtered PBS. Six 60 s videos with a camera level of 13 were obtained at room temperature using the NTA 2.2 software (Malvern Panalytical). The temperature for each run was logged before analysis. Videos were analysed using the NTA 2.2 software with a threshold of 10 and with a gain of 1.

### Transmission electron microscopy of EVs

Plasma-derived EVs were fixed with 4 % (v/v) paraformaldehyde and diluted 1 in 25 in 0.1 µm-filtered PBS. 5 µl of diluted EVs were applied to freshly glow-discharged carbon film copper grids (Agar Scientific, Essex, UK) for 3 min and then stained with two 5 µl drops of 1 % (w/v) uranyl acetate, before blotting with filter paper to dry. EVs were visualised using a JEOL JEM1400 transmission electron microscope (JEOL UK Ltd, Welwyn Garden City, UK) with an accelerating voltage of 120 kV. Images were captured using an EMSIS Xarosa digital camera (EMSIS GmbH, Münster, Germany) with Radius software.

### SDS-PAGE and immunoblot analysis

EV markers in plasma-derived EVs were examined by SDS-PAGE and immunoblot analysis as described previously^30^. Equal volumes of EV lysates or 20 µg of total cell lysates were separated by 10 % (v/v) SDS-PAGE and transferred onto nitrocellulose membranes.

Membranes were blocked with Tris-buffered saline Tween 20 (TBST) containing 5 % (w/v) low fat milk for 1 h and then and incubated overnight at 4°C with the following primary antibodies diluted in TBST-milk; mouse anti-CD81 (Thermo Fisher Scientific, Swindon, UK, 1:1500), rabbit anti-Tsg101 (Sigma, 1:1000), rabbit anti-calnexin (Cell Signalling Technology, Leiden, The Netherlands; 1:1000), or rabbit anti-ApoAI (R&D Systems, Abingdon, UK; 1:2000). Membranes were washed three times with TBST and incubated with an anti-rabbit-HRP conjugated secondary antibody (Vector Labs, Kirtlington, UK; 1 in 5000 dilution) or an anti-mouse-HRP antibody (Vector Labs; 1 in 5000 dilution) for 1 h. Membranes were washed three times with TBST and developed with Westar Supernova ECL detection reagent (Cyanagen, Bologna, Italy) and imaged using a G:BOX Chemi XX9 gel imaging system (Syngene, Cambridge, UK).

### Reverse Transcription Quantitative PCR (RT-qPCR) analysis of miRNAs in plasma-derived EVs

Total plasma-derived EVs eluted from the SEC columns (50 µl) were immediately lysed in 230 µl of lysis buffer C from the Maxwell RSC miRNA plasma and serum kit (Promega, Southampton, UK) supplemented with 1 µl of the RNA isolation spike-in control comprising pre-mixed UniSp2, UniSp4 and UniSp5 (339390, Qiagen, Manchester, UK) and incubated for 10 min on ice. miRNA was isolated from the EV lysates using the Maxwell RSC miRNA from plasma and serum kit (Promega) on a Maxwell RSC instrument (Promega) according to the manufacturer’s instructions and eluted in 40 µl of nuclease free water. RT-qPCR were performed as described previously^30^. RNA (4 µl) was reverse transcribed using the miRCURY LNA RT kit (Qiagen) together with the cDNA synthesis spike-in controls UniSp6 and cel-miR-39-3p. Samples were incubated at 42°C for 1 h, followed by inactivation of the RT enzyme at 95°C for 5 min. A master mix was prepared for each sample with 500 µl of miRCURY LNA miRNA SYBR Green master mix, 490 µl of water and 10 µl of cDNA. The master mix for each sample (10 µl) was pipetted into wells of the Human Serum/Plasma Small Focus (miScript) miRCURY LNA miRNA PCR panel (YAHS-206Z) (Qiagen). Real time quantification of the miRNAs was conducted on a LightCycler 480 instrument (Roche Diagnostics Limited, Burgess Hill, UK) with an initial activation at 95°C for 2 min followed by 45 cycles of 95°C for 10 s and 56°C for 1 min.

### Normalisation of RT-qPCR data

qPCR curves were analysed with the LightCycler 480 software (release 1.5.0) using the fit point method. In this method, a noise band is set to exclude background noise and crossing point (Cp) values are defined as the intersection between a horizontal threshold line and a log-line fitting the exponential portion of the amplification curve. The noise bands and the horizontal threshold lines were set separately for each miRNA in the panel and were kept the same across all samples.

Spike-in controls were used to assess the efficiencies of RNA isolation and cDNA synthesis for each sample. Thresholds for the five spike-in controls (UniSp2, UniSp4, UniSp5, UniSp6, cel-miR-39-3p) were calculated as follows: Cp values for each spike-in control were averaged across all samples. The standard deviation (SD) x2 of the average Cp values for each spike-in control across all samples was calculated. Samples with three or more spike-in controls with Cp values above or below these SD x2 thresholds indicated RNA isolation or cDNA synthesis efficiencies were outside the normal range and were removed from analysis.

NormFinder on R (version 4.5.0)^31^ was used to test the miRNAs from the miRCURY LNA miRNA PCR panel, allowing the identification of hsa-miR-16-5p and hsa-let-7a-5p as a pair of miRNAs with stable expression across our conditions. NormFinder allows identification of miRNAs that are not only stably expressed across the whole data set but are also stable between groups i.e., healthy controls and patients^31^. All Cp values were converted to absolute quantity using the 2^-ct^ formula. miRNA absolute quantities were then normalised against the geometric mean of the miRNAs selected for normalisation, hsa-miR-16-5p and hsa-let-7a-5p.

### Identification of significantly differentially expressed miRNAs

We only examined expression differences between miRNAs that had a minimum of three replicates per sex and condition, leaving a total of 63 miRNAs. All analyses were conducted in R (v.4.3.1). Normality of data for each miRNA was determined via a Shapiro Test and differences in expression were determined via linear models for normally distributed data and general linear models for Gamma distributed data, with condition (healthy or schizophrenic), sex and any interaction effect as factors. *Post hoc* testing for interaction effects between sex and condition was performed using a Tukey test implemented via the emmeans package (v.1.10.3). miRNA expression was considered significantly different if *P* < 0.05. All code is available at https://doi.org/10.5281/zenodo.18208502. Corrections for multiple comparisons were not applied due to the small sample size and exploratory nature of the study.

### Correlation of miRNA expression with clinical measures

Correlation of miRNA expression with clinical measures was limited to the miRNAs for which significant differential expression between patients and controls were found. In the full sample, Pearson’s correlation coefficients were used to examine the relationship between miRNA expression and age, and linear regression models were used to examine the relationship between miRNA expression and cognitive measures, with age, sex, and diagnosis as covariates. In the patient group only, linear regression models were used to examine the relationship of miRNA expression to total scores from the BPRS and BNSS, with age and sex as covariates. Linear regression was used to determine if miRNA expression was associated with white matter microstructure using age, sex, diagnosis and smoking status as covariates. Due to limited sample size, these analyses were considered exploratory and were not corrected for multiple comparisons. Analyses were performed using SPSS v.29.

### Gene ontology pathway enrichment analysis and STRING protein interaction network analysis of potential target genes

The miRNA database miRDB^32^ was used to identify putative target genes of the miRNAs significantly differentially expressed between conditions. Only target genes with a predicted target score threshold ≥80 were selected. Potential gene targets that were shared between 2 - 3 miRNAs were identified using Venn diagram analysis (https://bioinformatics.psb.ugent.be/webtools/Venn/). These shared gene targets were further analysed using gene ontology (GO) pathway enrichment analysis (PANTHER 19.0) (GO Ontology database DOI: 10.5281/zenodo.15066566, released 2025-03-16) with Fisher’s correction and only showing results with an FDR of *P* < 0.05. Protein clusters associated with specific biological functions were identified using the protein interaction network database STRING version 12.0^33^ with a high confidence score of 0.7 and using all sources of evidence.

### Diffusion tensor imaging

The imaging data were collected on a Siemens Prisma 3 Tesla scanner using a 64-channel coil. The data were collected using a multiband, echo-planar, spin-echo, T2-weighted sequence (TE/TR/Multiband Factor=97/4000ms/4 with the FOV=200 mm) with an isotropic spatial resolution of 1.6 mm. Diffusion data were preprocessed using HCP Diffusion pipeline^34,35^ that was combined with DESIGNER diffusion preprocessing tools^36^. Fractional anisotropy (FA) maps were obtained by fitting the diffusion tensor model using the FSL-FDT toolkit^37^. The population-based cerebral white matter tract atlas developed at John Hopkins University and distributed with the FSL package^38^ was used to calculate average FA values along the spatial course of major white matter tracts. The whole brain average FA was the primary measure. The data were processed using FSL’s tract-based spatial statistics (TBSS; http://fsl.fmrib.ox.ac.uk/fsl/fslwiki/TBSS) analytic method modified to project individual FA values onto the ENIGMA-DTI skeleton. DTI data were only available for 37 participants.

## Results

### Confirmation of the isolation of EVs from plasma

The size distributions of isolated plasma-derived EVs isolated were examined using NTA and revealed a mode diameter of 185 nm (Figure 1A). Negative staining of EVs followed by TEM confirmed the presence of EVs (Figure 1B). The majority of the EVs had a cup-shaped morphology which is typically seen in TEM preparations of EVs^39^. Immunoblot analysis of plasma-derived EVs confirmed the presence of the EV markers CD81 (Figure 1C) and Tsg101 (Figure 1D). As observed previously with EVs isolated from a separate cohort of healthy control plasma samples^30^, Tsg101 in plasma-derived EVs had a higher molecular weight than expected (Figure 1D), which may be due to post-translational modifications of proteins in EVs^40^. ApoA1 was detected in the EV preparations as before (Figure 1E)^30^, which is expected since lipoproteins are a similar size to EVs and therefore elute from the Exospin columns in the same fraction. The endoplasmic reticulum marker calnexin was absent from the EV preparations indicating enrichment of EVs without contamination from cell debris (Figure 1F).

**Figure 1.**
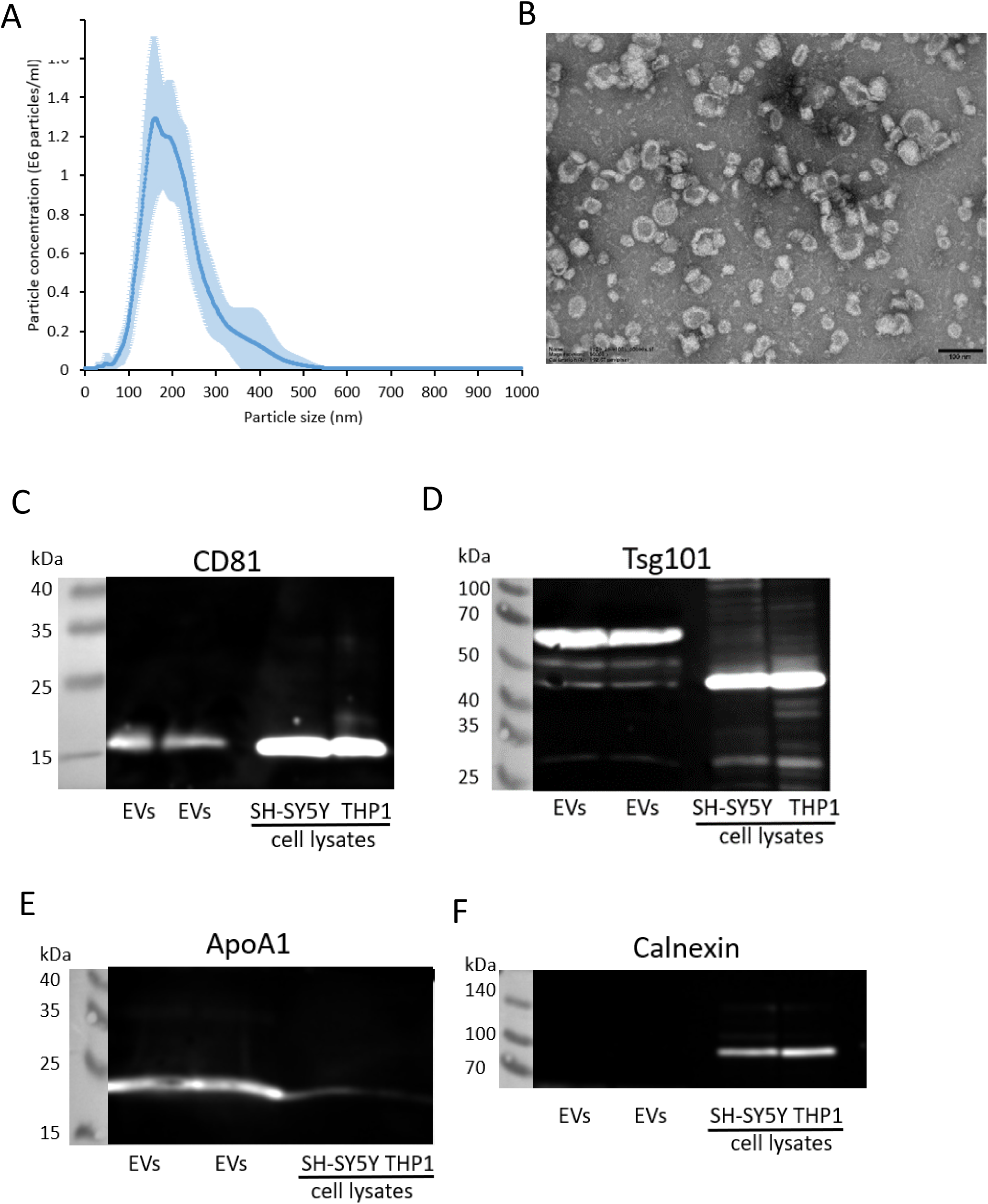
Characterisation of EVs isolated from plasma EVs were isolated from human plasma using size exclusion chromatography columns and examined using A) NTA, B) TEM (scale bar = 100 nm), and immunoblot analysis for CD81 (C), Tsg101 (D), ApoA1 (E), calnexin (F). Blots show two EV isolations from the plasma of the same healthy control sample. Cell lysates from the neuroblastoma cell line SH-SY5Y and monocytic cell line THP-1 were examined alongside.

### Eight miRNAs were significantly differentially expressed in plasma-derived EVs

RT-qPCR data from two individuals were removed from analysis due to three spike-in control values for these samples being outside the range of the set thresholds. Data from another two individuals were removed due to haemolysis. This resulted in 67 samples suitable for further analysis: 16 female healthy controls, 18 male healthy controls, 15 female schizophrenia patients, and 18 male schizophrenia patients. Supplemental Table 1 in Supplemental File 1 shows the characteristics of the 67 individuals with samples suitable for analysis.

The two most stable miRNAs for the normalisation of the data within the RT-qPCR panel were hsa-miR-26a-5p (stability score 0.08) and hsa-miR-126-3p (stability score 0.09) which had a paired stability score of 0.06. However, both of these miRNAs have been associated with schizophrenia in previous studies^7,8,41^. Therefore, the data were normalised against the geometric mean of the next two most stable miRNAs hsa-miR-16-5p (stability score 0.1) and hsa-let-7a-5p (stability score 0.1) which had a paired stability score of 0.07. Since the samples were analysed in two batches, batch effects were normalised as described in Supplemental File 1.

A total of eight out of 63 miRNAs were found to have a nominally significant difference either between conditions, sexes, or an interaction between sex and condition (Table 1). Two miRNAs were shown to have significantly higher expression in patients with schizophrenia compared to healthy controls; hsa-miR-103a-3p: X2 = 4.01, p = 0.045 (Figure 2A), hsa-miR-30e-5p: X2 = 13.4, p = 0.0002 (Figure 2B) and one had significantly lower expression; hsa-miR-200b-3p: X2 = 7.43, p = 0.006 (Figure 2C). A technical replicate for hsa-miR-103a-3p on the qPCR panel also showed a trend for increased expression in patients compared to controls (X2 = 2.91, p = 0.088, fold change = 1.5).

**Figure 2.**
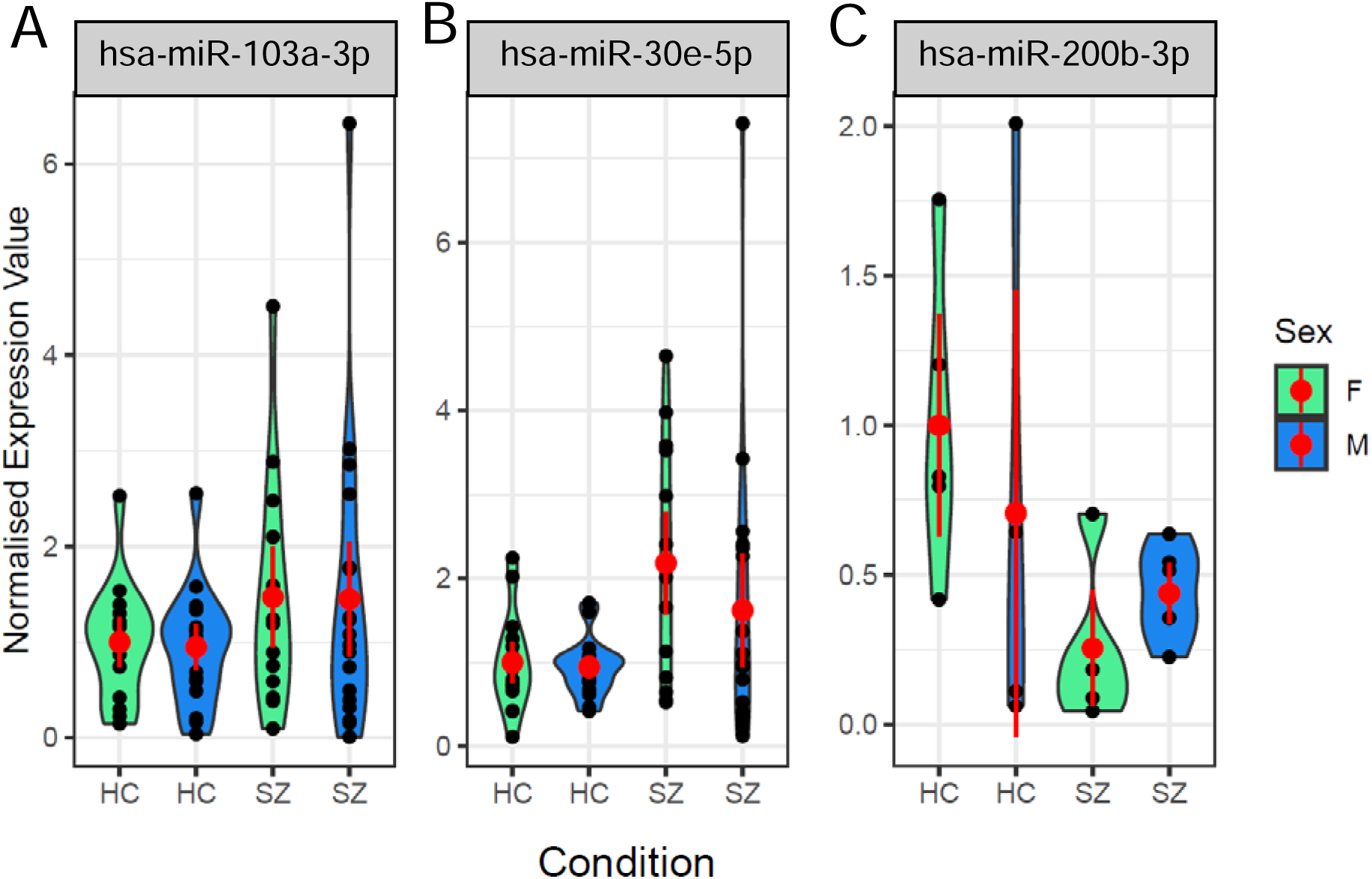
Three miRNAs were differentially expressed between controls and patients with schizophrenia miRNA expression in plasma-derived EVs was examined using RT-qPCR. The violin plots show the expression of three miRNAs significantly different between healthy controls and patients with schizophrenia: A) hsa-miR-103a-3p, B) hsa-miR-30e-5p, C) hsa-miR-200b-3p. Each black dot represents a replicate sample; the red dot represents the mean with 95% confidence intervals. HC = healthy control. SZ = schizophrenic patient. F = female. M = male.

**Table 1.** Fold changes and p-values of the eight significantly differentially expressed miRNAs Fold changes in miRNA expression were calculated from the averaged normalised expression values for each miRNA as all healthy controls vs. all schizophrenia patients for condition, all female healthy controls vs. all female schizophrenia patients for condition:sex interaction, and all females vs. all males for sex. *P*-values were derived from a linear model for hsa-miR-143-3p and hsa-miR-17-5p, and a general linear model with Gamma distribution for all others.

| **miRNA** | **Comparison** | **Fold change** | **p-value** | **F value** | **Chi squared value** |
| --- | --- | --- | --- | --- | --- |
| hsa-miR-103a-3p | Condition | 1.5 | 0.0452 | N/A | 4.01 |
| hsa-miR-200b-3p | Condition | -2.44 | 0.0064 | N/A | 7.43 |
| hsa-miR-30e-5p | Condition | 1.92 | 0.00025 | N/A | 13.4 |
| hsa-miR-143-3p | Condition:Sex | 2.26 | 0.021 | 5.77 | N/A |
| hsa-miR-423-5p | Sex | 2.17 | 0.0015 | N/A | 10.02 |
| hsa-miR-17-5p | Sex | -1.41 | 0.043 | 4.26 | N/A |
| hsa-miR-17-3p | Sex | -2.38 | 0.038 | N/A | 4.28 |
| hsa-miR-451a | Sex | -1.51 | 0.042 | N/A | 4.14 |

Only one miRNA, hsa-miR-143-3p, interacted between sex and condition and was significantly upregulated in females with schizophrenia compared to males with schizophrenia and healthy controls (linear model, F = 5.77, p = 0.021) (Supplemental Figure 1 in Supplemental File 1). We also identified four miRNAs that were differentially expressed between sexes (Supplemental Figure 2 in Supplemental File 1). Three showed reduced expression in females compared to males, hsa-miR-17-3p (X2 = 4.28, p = 0.038), hsa-miR-451a (X2 = 4.14, p = 0.042) and hsa-miR-17-5p (F = 4.26, p = 0.043), although for hsa-miR-17-5p, it is clear the effect is driven by healthy control females (Supplemental Figure 2 in Supplemental File 1). In contrast, hsa-miR-423-5p showed higher expression in females compared to males (X2 = 10.02, p = 0.0015).

### Putative gene targets of miRNAs differentially expressed in schizophrenia are involved in biological functions associated with neurogenesis

First, the miRNA database miRDB was used to identify putative gene targets with a predicted target score of ≥80 for the three miRNAs differentially expressed between conditions. Venn diagram analysis revealed 136 genes that had the potential to be regulated by combinations of 2 - 3 of these miRNAs (Figure 3A and Supplemental Table 2 in Supplemental File 1). Only three potential target genes were shared between all three miRNAs: sodium voltage-gated channel alpha subunit 8 (*SCN8A*), Janus kinase and microtubule interacting protein 2 (*JAKMIP2*), and the guanine nucleotide exchange factor DENN domain containing 1B (*DENND1B*).

**Figure 3.**
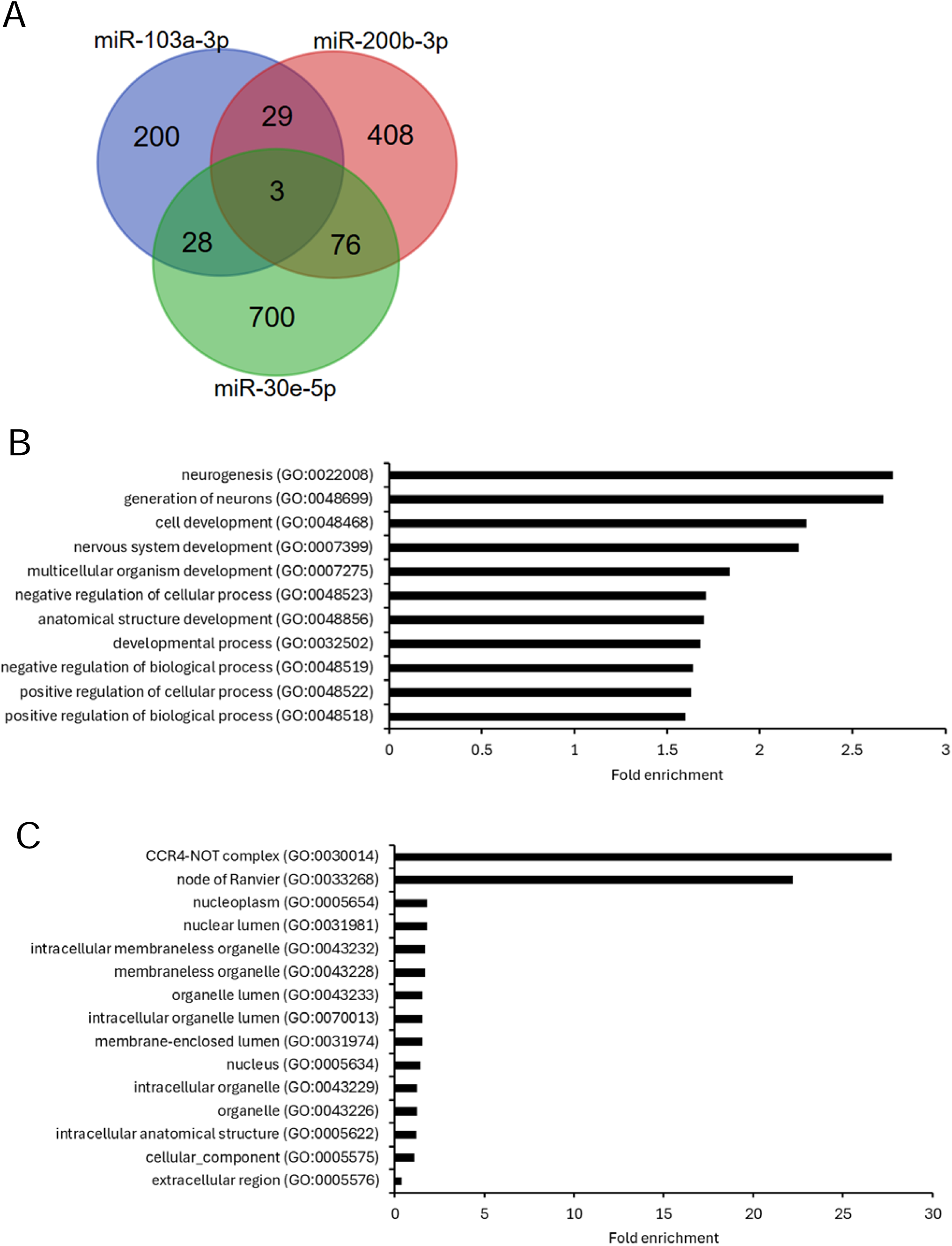
Identification of shared potential gene targets of the 2-3 differentially expressed miRNAs and gene ontology enrichment analysis Putative target genes of the three miRNAs significantly differentially expressed between patients and controls were identified using miRDB. A) Venn diagram analysis identified 136 potential target genes that were shared between 2 - 3 of these miRNAs. The 136 shared target genes were analysed using gene ontology enrichment analysis and revealed enrichment of biological process terms (B) and cellular component terms (C) associated with these gene targets. FDR p<0.05.

We next performed gene ontology enrichment analysis to discover groups of target genes associated with similar biological and cellular processes. GO term enrichment analysis of the 136 shared target genes revealed enrichment of biological process terms associated with the nervous system and development including neurogenesis, cell development, nervous system development, and regulation of cellular processes (Figure 3B, Supplemental Table 3A in Supplemental File 1). Gene targets were also enriched for the cellular component terms CCR4-NOT complex and node of Ranvier (Figure 3C, Supplemental Table 3B). Lists of genes associated with enriched GO biological process terms and cellular component terms for gene targets shared between 2 - 3 miRNAs are shown in Supplemental File 2.

STRING protein interaction network analysis was performed to examine potential interactions between the identified target genes. This analysis found that the protein interaction network derived from the 136 putative target genes included 22 edges compared to the expected 8 edges (*P* = 3.45 × 10^-5^), indicating that the clustering of potential target genes into similar biological functions did not occur by chance. Target genes were enriched in clusters involved in circadian rhythm (*CLOCK, RORA, ATXN1*), the node of Ranvier (*SCN8A, ANK3, SCN2A*), the CCR4-NOT complex which regulates gene expression^42^ (*CNOT9, CNOT6, CPEB3*), and the FAR/SIN/STRIPAK complex which are scaffolding proteins that regulate signalling pathways^43^ (*STRN, PDCD10, MAP4K4*) (Figure 4A).

**Figure 4.**
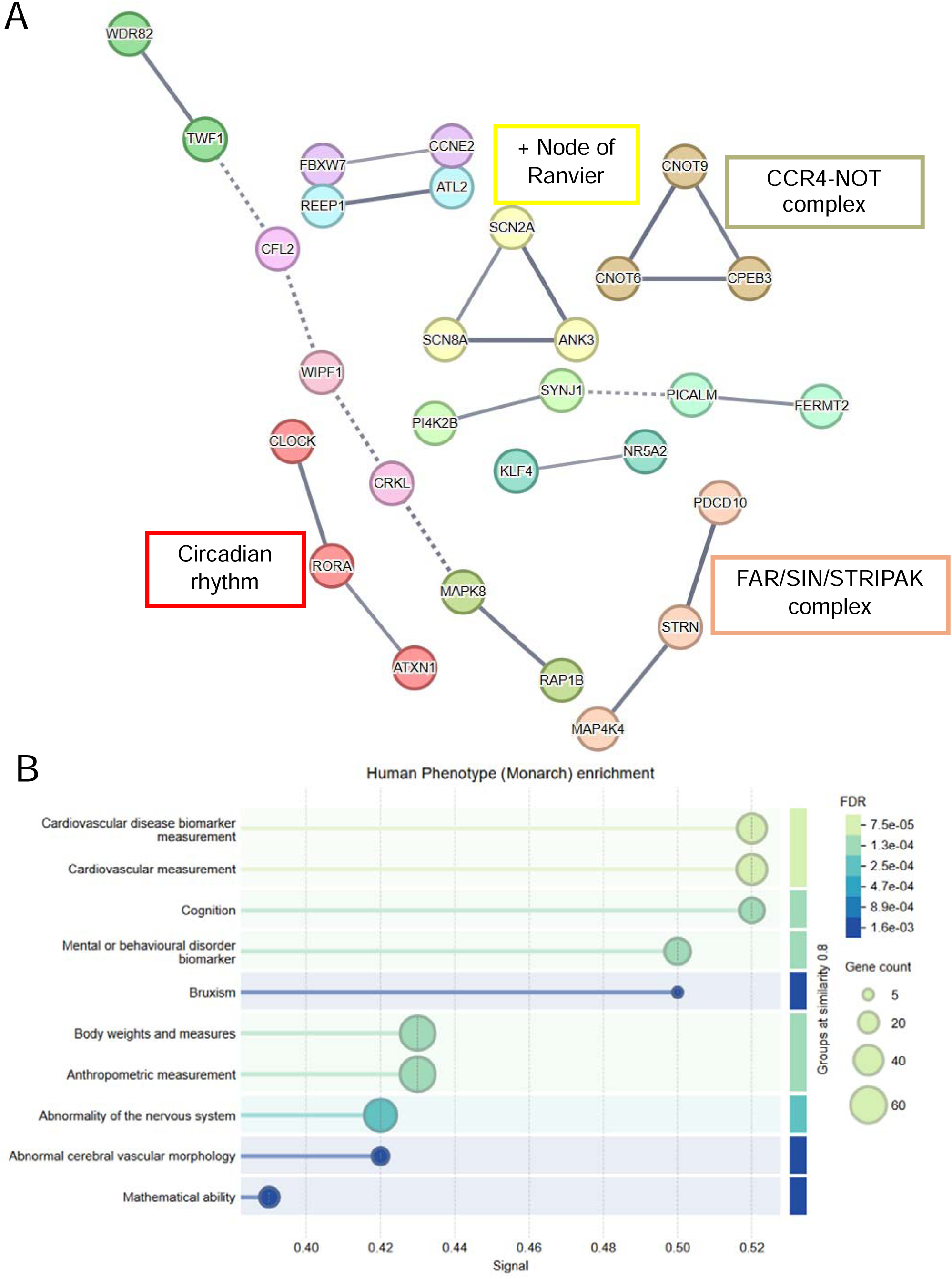
STRING protein interaction network analysis of shared potential gene targets A) The 136 potential target genes shared between 2-3 miRNAs differentially expressed between patients and controls were analysed using STRING protein interaction network analysis with a high confidence score of 0.7 and network edges showing confidence strength. B) Human phenotype (Monarch) enrichment using STRING.

Monarch human phenotype enrichment analysis showed enrichment of phenotypes related to cardiovascular disease, cognition and mental/behavioural disorder biomarker (Figure 4B).

### Correlation of miRNA expression with clinical and imaging measures

In the cohort of 67 individuals tested, age was significantly correlated with expression of hsa-miR-103a-3p (r = 0.30, p = 0.015), but not with hsa-miR-200b-3p or hsa-miR-30e-5p. This aging effect was driven by patients (r = 0.48, p = 0.006) and was not evident in controls (r = 0.08, p = 0.67). Additionally, in patients hsa-miR-30e-5p was significantly correlated with age (r = 0.48, p = 0.005) but this was not found in controls (r = -0.17, p = 0.35). In the combined sample, expression of hsa-miR-103a-3p was significantly correlated with working memory (β = 0.28, p = 0.032), but not with processing speed (β = 0.16, p = 0.21). The correlation between hsa-miR-103a-3p and working memory was positive but not significant in either patients (β = 0.25, p = 0.23) or controls (β = 0.35, p = 0.073). No significant correlations were found between hsa-miR-200b-3p or hsa-miR-30e-5p and cognitive measures. In the patient group of 33 individuals, no miRNA expression alterations were significantly associated with symptom measures (BPRS or BNSS scores).

Whole brain tract-averaged FA was next analysed in a cohort of 37 participants including patients and controls to explore a possible relationship between miRNA expression and white matter microstructure. We found that this metric negatively correlated with hsa-miR-103a-3p expression in the whole sample at a trend level (β = -0.34, p = 0.053). As smoking may influence FA levels^44^, we repeated this analysis including smoking as a covariate, and in this analysis the association was significant (β = -0.41, p = 0.02), with similar but non-significant trends for patients (β = -0.41, p = 0.10) and controls (β = -0.15, p = 0.63) when examined separately. White matter FA was not significantly associated with hsa-miR-200b-3p or hsa-miR-30e-5p.

## Discussion

Previous studies have shown that miRNA expression is altered in the CNS^11,12,13,14,15^ and peripheral circulation^17,18,19,20,21^ of patients with schizophrenia. Changes in miRNA expression levels in patients may have functional consequences due to the role of these miRNAs in regulating processes essential for nervous system development and function such as synaptogenesis and transcriptional regulation^10^. To date, a limited number of studies have examined miRNA expression in blood-derived EVs in schizophrenia patients compared to healthy controls^7,8,22,23,45,46^. In this study, we isolated EVs from plasma samples to examine changes in miRNA levels compared to healthy controls in an attempt to identify a “miRNA signature” in patients with schizophrenia and correlate miRNA expression with clinical measures relevant to schizophrenia.

We identified three significantly differentially expressed miRNAs in EVs isolated from patients with schizophrenia compared to healthy controls; upregulation of hsa-miR-103a-3p and hsa-miR-30e-5p, and downregulation of hsa-miR-200b-3p. Our data confirm previous studies that have shown increased expression of miR-103a-3p in the peripheral circulation of patients with schizophrenia. Du *et al*. (2023)^7^ detected significant upregulation of miR-103a-3p in plasma-derived EVs from patients with first episode schizophrenia compared to healthy controls in a small sample of individuals. A recent study by Chen *et al.* (2025)^45^ also detected significantly increased expression of miR-103a-3p in blood-derived EVs from patients with SZ, further validating our findings. Jeffries *et al.* (2016)^47^ detected altered expression of miR-103a-3p in PBMCs derived from patients who later developed psychosis compared to patients who did not progress to psychosis, and a follow-on study using the same samples associated miR-103a-3p with cortical thinning in patients who later developed psychosis^48^.

Several previous studies have shown increased miR-30e-5p expression in the plasma^19, 49,50^ and PBMCs^19,51^ of patients with schizophrenia compared to healthy controls. We have shown for the first time that levels of miR-30e-5p are also significantly increased in plasma-derived EVs from patients with schizophrenia. Sun *et al*. (2015)^19^ suggested that increased miR-30e-5p expression in plasma from patients with schizophrenia could have a diagnostic use as a biomarker of schizophrenia. Furthermore, overexpression of miR-30e-5p in the hippocampus of rats resulted in cognitive deficits in these animals^52^, indicating that the observed increases in miR-30e-5p expression in the peripheral circulation may underly functional roles of this miRNA in schizophrenia. Finally, hsa-miR-200b-3p was significantly downregulated in patients with schizophrenia compared to healthy controls. This miRNA has been shown to target the expression of N-methyl D-aspartate (NMDA) receptor subunits resulting in cognitive impairments in rats^53^.

Next, we identified putative target genes of the three miRNAs differentially expressed between conditions. GO term enrichment analysis of the putative target genes that were shared between 2 - 3 of the significantly differentially expressed miRNAs revealed enrichment of several biological process terms associated with development of the nervous system such as neurogenesis, cell development and regulation of cellular processes. This is in agreement with a previous study by Barnett *et al*. (2023)^8^ that identified target genes of miRNAs differentially expressed in neuronally-derived EVs from patients with schizophrenia and found enrichment of GO terms related to synapse organisation and regulation of synapse structure. It is possible that aberrant expression of the three miRNAs found to be differentially expression in patients in our study reflects disruption in neurogenesis and nervous system development that may contribute to schizophrenia in susceptible individuals, although further investigation into the role of these miRNAs in neuronal development is required. STRING protein interaction network analysis also revealed enrichment of target genes associated with circadian rhythm. This is relevant because patients with schizophrenia often have disrupted sleep^54^. A study by Johansson *et al.* (2016)^55^ investigated whether patients with schizophrenia have differences in the molecular clock which regulates the sleep-wake cycle and showed that PBMCs isolated from patients with first episode psychosis had reduced expression of several proteins involved in regulating circadian rhythm, including the main regulator of circadian rhythm, Circadian Locomotor Output Cycles Kaput (CLOCK)^55^. Since CLOCK was one of the putative target genes of the miRNAs significantly and differentially expressed in our study, it would be relevant to further investigate whether these miRNAs can regulate the expression of circadian rhythm proteins such as CLOCK in neurons.

Both GO enrichment analysis and STRING protein interaction network analysis of potential target genes identified a protein cluster associated with the node of Ranvier. This is of interest because nodes of Ranvier are required for propagation of action potentials along neurons for rapid neurotransmission and it has been proposed that there are abnormalities in these structures in patients with schizophrenia^56^. Nodes of Ranvier are enriched in voltage-gated sodium channels and cytoskeletal proteins such as ankyrin-3 (ANK3), both of which were identified as potential target genes of the three differentially expressed miRNAs in our study. Ankyrin-3 is a large scaffolding protein which organises node structure and stabilises clusters of voltage-gated sodium channels within the nodes^57,58^. Interestingly, significantly reduced expression of both the voltage-gated sodium channel SCN8A and ankyrin-3 were detected in post-mortem brain tissue of patients with schizophrenia compared to controls^59^. Roussos *et al*. (2012)^59^ also associated an *ANK3* variant with reduced ankyrin-3 expression and cognitive deficits, suggesting that downregulation of *ANK3* may have functional consequences for neurotransmission in patients with schizophrenia. It would thus be worthwhile examining whether the three differentially expressed miRNAs, either alone or in combination, can directly affect neurogenesis, circadian rhythm or neurotransmission by regulating the expression of these predicted target genes.

To examine potential clinical associations between the three significantly differentially expressed miRNAs observed in our study, expression of these miRNAs was correlated with clinical measures including cognition and white matter integrity. Correlation analysis of combined data from controls and patients revealed that miR-103a-3p expression was associated with working memory and negatively correlated with white matter integrity, but these correlations were not specific to patients. A previous study associated miR-103a-3p expression in leukocytes with grey matter changes and cortical thinning in patients who later developed psychosis^48^. Another study detected higher expression of miR-103a-3p in serum-derived exosomes from patients with MDD compared to controls and found that overexpression of this miRNA reduced the differentiation of rat neuronal cells through the downregulation of brain-derived neurotrophic factor (BDNF)^60^. The authors also showed that inhibition of miR-103a-3p by intra-cerebroventricular injection of a miR-103a-3p antagomiR reduced depressive behaviours in a rat model of chronic unpredictable mild stress^60^, further implicating this miRNA in impaired neuronal development. Barnett *et al*. (2023)^8^ used small RNA sequencing to examine the expression of miRNAs in serum-derived EVs of neuronal origin and correlated expression of several miRNAs with cognitive deficits in patients with schizophrenia. The authors observed significant differences in miRNA expression between patients and controls, which were more pronounced in patients with cognitive deficits but did not identify any of the miRNAs found to be significantly differentially expressed in our study. This may be due to differences in the source of the EVs (neuronal compared to total plasma EVs) or different patient cohorts. It is important to consider that the expression of the three miRNAs identified in our study did not correlate with symptom severity. Furthermore, the finding that miR-103a-3p was associated with working memory and white matter integrity across the whole sample rather than being specific to patients potentially indicates expression of this miRNA may reflect individuals with an increased susceptibility to developing schizophrenia rather than being associated with diagnosis.

A potential limitation of this study is the modest number of patients and controls examined, especially for the imaging analysis where data for only 37 individuals were available. Larger sample sizes, preferably from separate cohorts would help to validate our findings. We also focused solely on patients with schizophrenia, but it would be interesting to compare miRNA signatures in plasma-derived EVs from patients with other psychiatric conditions with similar symptoms such as bipolar disorder, to determine whether the observed changes in miRNA expression are specific to schizophrenia or generally related to psychosis. For example, Smalheiser *et al*. (2014)^15^ found that there was an overlap in the expression levels of several significantly differentially expressed miRNAs in the PFC of patients with schizophrenia and bipolar disorder, whereas as other miRNAs were specific to each condition. The two miRNAs identified as significantly upregulated in plasma-derived EVs from schizophrenia patients in our study, miR-103a-3p and miR-30e-5p, have been shown to have increased expression in schizophrenia in previous studies as described above. However, studies examining miRNA expression in the peripheral circulation of patients with schizophrenia have produced variable results, and we did not detect changes in expression of all miRNAs previously associated with schizophrenia. This may be due to different EV isolation protocols, the various methods used for miRNA quantification such as RT-qPCR, microarrays and RNA sequencing, different patient cohorts, and the heterogeneity of schizophrenia. It is likely that a wider range of differentially expressed miRNAs could have been detected using a more sensitive approach such as small RNA sequencing, and this may explain differences between our results and previous studies. However, the RT-qPCR panel used here was specifically for plasma/serum-enriched miRNAs and included several miRNAs previously associated with schizophrenia such as miR-206^22^ and miR-223-3p^61^. Drug treatment may also affect miRNA levels in patients. Du *et al*. (2019)^22^ found that miR-206 was significantly upregulated in first-episode drug-free patients but not in patients on chronic long-term medication. Furthermore, *in vitro* experiments showed that the anti-psychotic clozapine increased miR-675-3p expression in the neuroblastoma cell line SH-SY5Y^23^, highlighting the potential influence of medication on miRNA levels.

In total, this study identified a miRNA signature of three differentially expressed miRNAs in plasma-derived EVs from patients with schizophrenia and for the first time showed significantly increased expression of miR-30e-5p in plasma-derived EVs from patients.

Furthermore, shared predicted target genes of these three miRNAs involved in neurogenesis and circadian rhythm together with the correlation of miR-103a-3p expression with working memory and white matter integrity may underlie the importance of altered expression of these miRNAs in nervous system development and may indicate their involvement in the development of schizophrenia alongside other genetic and environmental factors.

## Acknowledgements

We thank all blood donors and phlebotomists who took part in this study.

## Ethical approval

This study was approved by the Institutional Review Board of the University of Maryland Baltimore. All participants provided informed written consent.

Ethical approval was obtained from the University of Leicester Ethical Practices Committee (application 24427-fg36-ls:genetics&genomebiology).

