## Supplemental File 1 for "Identification of a miRNA signature for schizophrenia in plasma-derived extracellular vesicles"

**Identification of a miRNA signature for schizophrenia in plasma-derived extracellular vesicles**

Mary E W Collier^1*^, Joshua Chiappelli^2^, Hollie Marshall^1^, Nicolas Sylvius^3^, Natalie Allcock^4^, Josh Whittingham^4^, Peter Kochunov^5^, Robert Schwarcz^2^, Elliot Hong^5^, Flaviano Giorgini^1^

^1^Division of Genetics and Genome Biology, University Road, University of Leicester, Leicester, UK, LE1 7RH

^2^Department of Psychiatry, Maryland Psychiatric Research Center, University of Maryland School of Medicine, Baltimore, Maryland, USA

^3^Genomics Core Facility, University of Leicester, Leicester, UK

^4^Electron Microscopy Facility, Hodgkin Building, Lancaster Road, University of Leicester, UK, LE1 7HG

^5^Department of Psychiatry and Behavioural Sciences, McGovern Medical School, Houston, Texas, USA

**Supplemental Methods**

**Removal of batch effects**

As data were generated in two batches, we normalized by batch by dividing the expression value for each miRNA by the mean expression of that miRNA for healthy control females. Whilst this is standard in the field, to ensure there were no overall sex/condition effects within the data, we calculated the coefficient of variation for each miRNA per sex/condition group per batch in R (v.4.3.1). A linear model showed that the variation within the data is not determined by sex (general linear model, X2 = 1.87, p = 0.17) or condition (general linear model, X2 = 0.26, p = 0.61), as such we tested normalizing by healthy control males and females, finding that using the female data reduced the batch effect to the greatest extent (general liner model, batch effect, male normalization: X2 = 4.41, p = 0.036, female normalization: X2 = 3.99, p = 0.046).

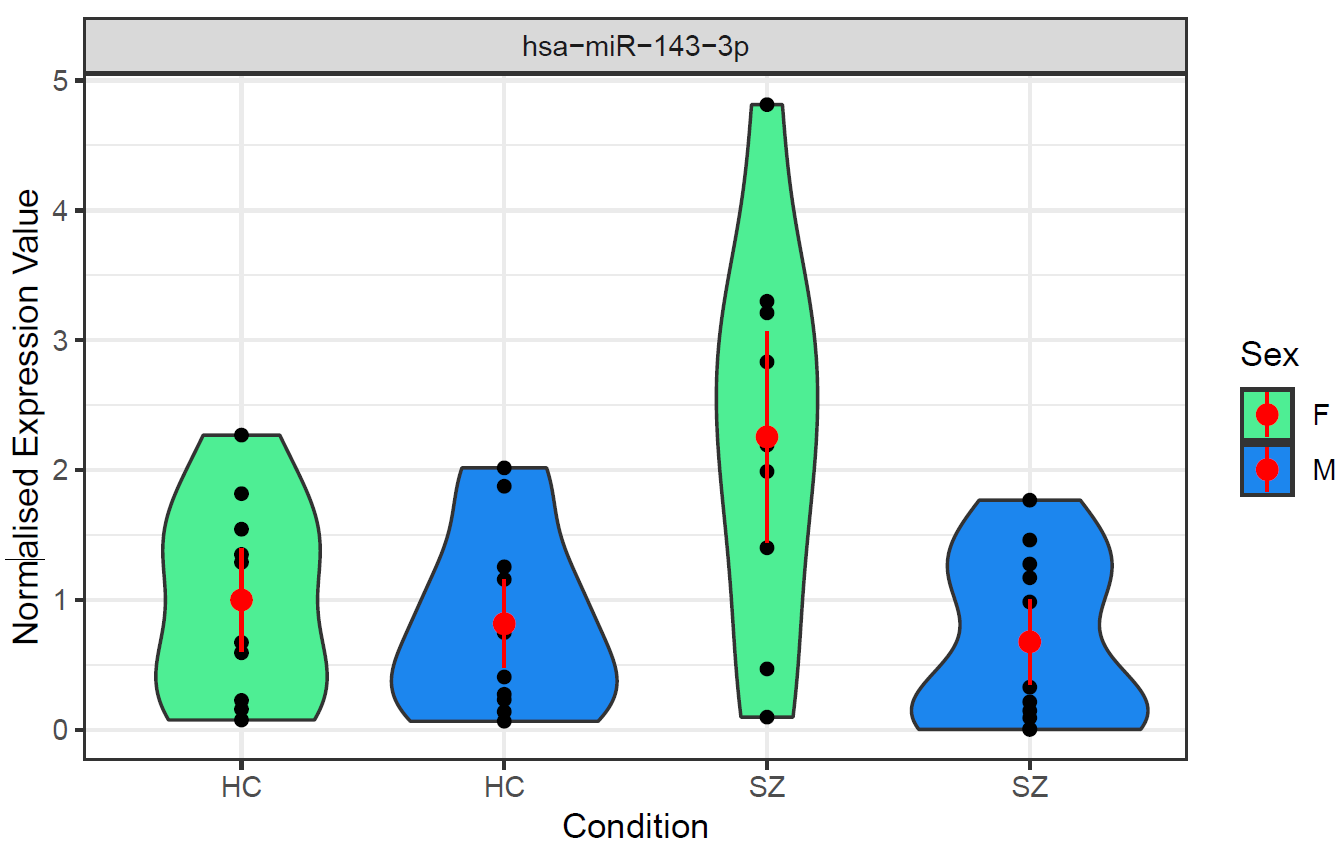

**Supplemental Figure 1.** Violin plot for hsa-miR-143-3p expression which showed a significant interaction between sex and condition. Each black dot represents a replicate sample; the red dot represents the mean with 95% confidence intervals. HC = healthy control. SZ = schizophrenic patient. F = female. M = male.

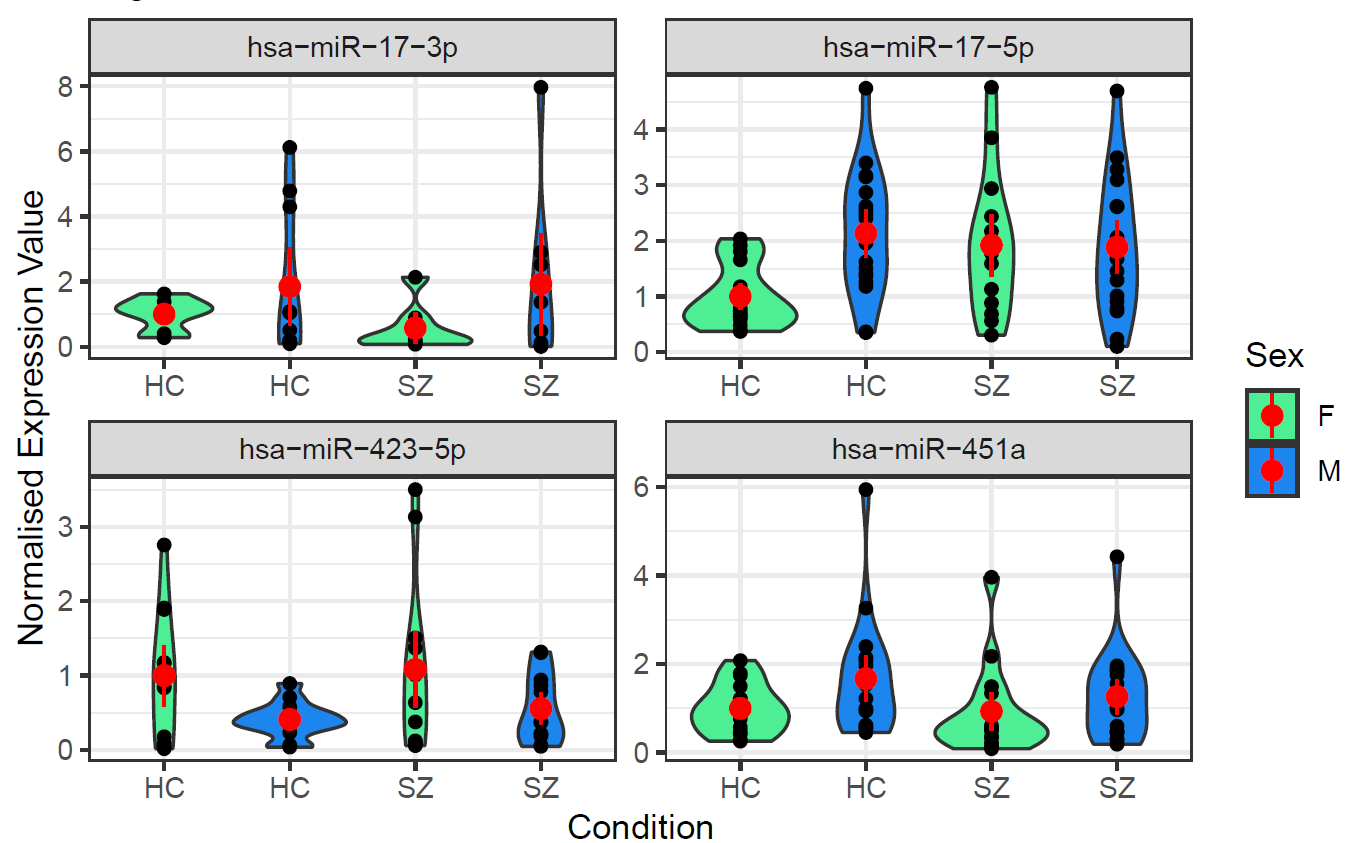

**Supplemental Figure 2**. Violin plots for miRNAs significantly differentially expressed between females and males. Each black dot represents a replicate sample. The red dot represents the mean with 95% confidence intervals. HC = healthy control. SZ = schizophrenic patient. F = female. M = male.

|  | **Patients (n=33)** | **Controls (n=34)** | **Test statistic** | **p-value** |
| --- | --- | --- | --- | --- |
| **Age** | 30.6 ± 8.5 | 31.4 ± 9.1 | t=0.49 | 0.62 |
| **Sex (M/F)** | 18 / 15 | 18 / 16 | χ2=0.02 | 0.90 |
| **Smoker/non-smoker** | 11/22 | 9/25 | χ2=0.38 | 0.54 |

**Supplemental Table 1.** Characteristics of the 67 individuals with samples suitable for analysis

| **miRNAs** | **Total number of shared gene targets** | **Gene target** |
| --- | --- | --- |
| miR-103a-3p; miR-200b-3p; miR-30e-5p | 3 | JAKMIP2 |
|  |  | DENND1B |
|  |  | SCN8A |
| miR-103a-3p; miR-200b-3p | 29 | AGFG1 |
|  |  | NFIA |
|  |  | PAG1 |
|  |  | RAB11FIP2 |
|  |  | ARIH1 |
|  |  | KLF4 |
|  |  | ZFPM2 |
|  |  | GPATCH8 |
|  |  | CCNYL1 |
|  |  | PPM1E |
|  |  | ELK4 |
|  |  | FERMT2 |
|  |  | KDM7A |
|  |  | UBE2R2 |
|  |  | PRKG1 |
|  |  | CEP85L |
|  |  | SRGAP1 |
|  |  | ZBTB10 |
|  |  | CAB39 |
|  |  | FBXW7 |
|  |  | ZDHHC21 |
|  |  | CSNK1G3 |
|  |  | SLITRK1 |
|  |  | SYNJ1 |
|  |  | NOVA1 |
|  |  | ANK3 |
|  |  | CLIP1 |
|  |  | STRN |
|  |  | GIT2 |
| miR-103a-3p; miR-30e-5p | 28 | GSKIP |
|  |  | HIC2 |
|  |  | CPEB3 |
|  |  | TWF1 |
|  |  | GPCPD1 |
|  |  | RAI14 |
|  |  | INO80D |
|  |  | DCUN1D3 |
|  |  | CCDC6 |
|  |  | NF1 |
|  |  | GALNT7 |
|  |  | PDCD10 |
|  |  | TNRC6B |
|  |  | CLOCK |
|  |  | KATNBL1 |
|  |  | WASHC4 |
|  |  | RORA |
|  |  | TDG |
|  |  | SCN2A |
|  |  | PHACTR2 |
|  |  | MAPK8 |
|  |  | SIX4 |
|  |  | ZCCHC2 |
|  |  | DLG5 |
|  |  | NSG1 |
|  |  | YTHDC1 |
|  |  | ROR1 |
|  |  | NDEL1 |
| miR-200b-3p; miR-30e-5p | 76 | RTKN2 |
|  |  | NUFIP2 |
|  |  | ERG |
|  |  | SNX16 |
|  |  | ATL2 |
|  |  | A1CF |
|  |  | CECR2 |
|  |  | CEP350 |
|  |  | DGKH |
|  |  | VAT1L |
|  |  | OTUD4 |
|  |  | PPP1R9A |
|  |  | KIAA0355 |
|  |  | PICALM |
|  |  | CNOT6 |
|  |  | HDAC9 |
|  |  | FAM126B |
|  |  | ZNF711 |
|  |  | GOLGA1 |
|  |  | NR3C1 |
|  |  | FRMD6 |
|  |  | ELAVL2 |
|  |  | BDP1 |
|  |  | OXR1 |
|  |  | PDS5B |
|  |  | COL4A3BP |
|  |  | DPY19L3 |
|  |  | OSTM1 |
|  |  | SLC4A7 |
|  |  | BCL11B |
|  |  | ELMOD2 |
|  |  | FIGN |
|  |  | WDR82 |
|  |  | CDH20 |
|  |  | REEP1 |
|  |  | PRDM1 |
|  |  | FOXG1 |
|  |  | CNOT9 |
|  |  | GLCCI1 |
|  |  | SEC23A |
|  |  | RAP1B |
|  |  | CFL2 |
|  |  | RASA2 |
|  |  | TBL1XR1 |
|  |  | NR5A2 |
|  |  | CCNE2 |
|  |  | PI4K2B |
|  |  | TOGARAM1 |
|  |  | ZEB2 |
|  |  | SLC25A36 |
|  |  | MBNL3 |
|  |  | CEP41 |
|  |  | ADAMTS3 |
|  |  | BRWD3 |
|  |  | LOX |
|  |  | KCTD8 |
|  |  | MMD |
|  |  | ATXN1 |
|  |  | S100PBP |
|  |  | MARCH6 |
|  |  | RAP2C |
|  |  | SIX1 |
|  |  | WIPF1 |
|  |  | IGF2R |
|  |  | ELL2 |
|  |  | MAP4K4 |
|  |  | PLPPR4 |
|  |  | PPP1R18 |
|  |  | ERRFI1 |
|  |  | BNC2 |
|  |  | CRKL |
|  |  | PHTF2 |
|  |  | EDEM3 |
|  |  | DPY19L1 |
|  |  | YPEL2 |
|  |  | SLC35B4 |

**Supplemental Table 2. Potential gene targets shared between 2-3 miRNAs**

Putative gene targets of the three miRNAs significantly differentially expressed between conditions were identified using miRDB. The table shows gene targets shared between 2 - 3 miRNAs which were identified using Venn diagram analysis.

| **A) GO biological process term** | **Number of genes in *Homo sapiens* reference list** | **Number of genes in uploaded data set that map to reference list** | **Number of genes in uploaded data set expected to map to reference list** | **Fold Enrichment of genes in data set compared to the expected number of genes** | **Raw P-value** | **False Discovery Rate** |
| --- | --- | --- | --- | --- | --- | --- |
| neurogenesis (GO:0022008) | 1413 | 26 | 9.54 | 2.72 | 2.54E-06 | 1.88E-02 |
| generation of neurons (GO:0048699) | 1218 | 22 | 8.23 | 2.67 | 2.24E-05 | 3.32E-02 |
| cell development (GO:0048468) | 2307 | 35 | 15.58 | 2.25 | 4.15E-06 | 1.54E-02 |
| nervous system development (GO:0007399) | 2279 | 34 | 15.39 | 2.21 | 7.66E-06 | 1.62E-02 |
| multicellular organism development (GO:0007275) | 4029 | 50 | 27.21 | 1.84 | 7.47E-06 | 1.85E-02 |
| negative regulation of cellular process (GO:0048523) | 5027 | 58 | 33.95 | 1.71 | 6.49E-06 | 1.92E-02 |
| anatomical structure development (GO:0048856) | 5304 | 61 | 35.82 | 1.7 | 3.53E-06 | 1.74E-02 |
| developmental process (GO:0032502) | 5830 | 66 | 39.38 | 1.68 | 1.67E-06 | 2.47E-02 |
| negative regulation of biological process (GO:0048519) | 5248 | 58 | 35.45 | 1.64 | 3.28E-05 | 4.41E-02 |
| positive regulation of cellular process (GO:0048522) | 5813 | 64 | 39.26 | 1.63 | 1.04E-05 | 1.71E-02 |
| positive regulation of biological process (GO:0048518) | 6113 | 66 | 41.29 | 1.6 | 9.76E-06 | 1.81E-02 |

| **B) GO cellular component term** | **Number of genes in *Homo sapiens* reference list** | **Number of genes in uploaded data set that map to reference list** | **Number of genes in uploaded data set expected to map to reference list** | **Fold Enrichment of genes in data set compared to the expected number of genes** | **Raw P-value** | **False Discovery Rate** |
| --- | --- | --- | --- | --- | --- | --- |
| CCR4-NOT complex (GO:0030014) | 16 | 3 | 0.11 | 27.76 | 1.58E-04 | 2.43E-02 |
| node of Ranvier (GO:0033268) | 20 | 3 | 0.14 | 22.21 | 3.16E-04 | 3.95E-02 |
| nucleoplasm (GO:0005654) | 4223 | 52 | 28.52 | 1.82 | 4.12E-06 | 4.12E-03 |
| nuclear lumen (GO:0031981) | 4596 | 56 | 31.04 | 1.8 | 1.81E-06 | 3.61E-03 |
| intracellular membraneless organelle (GO:0043232) | 5463 | 62 | 36.9 | 1.68 | 4.41E-06 | 2.94E-03 |
| membraneless organelle (GO:0043228) | 5463 | 62 | 36.9 | 1.68 | 4.41E-06 | 2.20E-03 |
| organelle lumen (GO:0043233) | 5759 | 60 | 38.9 | 1.54 | 1.30E-04 | 2.60E-02 |
| intracellular organelle lumen (GO:0070013) | 5759 | 60 | 38.9 | 1.54 | 1.30E-04 | 2.36E-02 |
| membrane-enclosed lumen (GO:0031974) | 5759 | 60 | 38.9 | 1.54 | 1.30E-04 | 2.16E-02 |
| nucleus (GO:0005634) | 7736 | 75 | 52.25 | 1.44 | 9.73E-05 | 2.16E-02 |
| intracellular organelle (GO:0043229) | 13488 | 115 | 91.1 | 1.26 | 9.04E-06 | 2.58E-03 |
| organelle (GO:0043226) | 14293 | 120 | 96.54 | 1.24 | 4.62E-06 | 1.85E-03 |
| intracellular anatomical structure (GO:0005622) | 15119 | 124 | 102.12 | 1.21 | 6.98E-06 | 2.32E-03 |
| cellular_component (GO:0005575) | 18937 | 138 | 127.9 | 1.08 | 2.19E-04 | 2.92E-02 |
| extracellular region (GO:0005576) | 4305 | 11 | 29.08 | 0.38 | 5.55E-05 | 1.39E-02 |

**Supplemental Table 3. GO term enrichment analysis of the 136 potential gene targets shared between the 2-3 miRNAs significantly differentially expressed between conditions.** Lists of genes associated with A) biological process terms and B) cellular component terms enriched in GO analysis of the 136 putative target genes shared between 2 - 3 miRNAs.
